# Identifying incident dementia by applying machine learning to a very large administrative claims dataset

**DOI:** 10.1101/396127

**Authors:** Vijay S. Nori, Christopher A. Hane, David C. Martin, Alexander D. Kravetz, Darshak M. Sanghavi

## Abstract

**INTRODUCTION:** Alzheimer’s disease and related dementias (ADRD) are highly prevalent conditions, and prior efforts to develop predictive models have relied on demographic and clinical risk factors using traditional logistical regression methods. We hypothesized that machine-learning algorithms using administrative claims data may represent a novel approach to predicting ADRD.

**METHODS:** Using a national de-identified dataset of more than 125 million patients including over 10,000 clinical, pharmaceutical, and demographic variables, we developed a cohort to train a machine learning model to predict ADRD 4-5 years in advance.

**RESULTS:** The Lasso algorithm selected a 50-variable model with an area under the curve (AUC) of 0.693. Top diagnosis codes in the model were memory loss (780.93), Parkinson’s disease (332.0), mild cognitive impairment (331.83) and bipolar disorder (296.80), and top pharmacy codes were psychoactive drugs.

**DISCUSSION:** Machine learning algorithms can rapidly develop predictive models for ADRD with massive datasets, without requiring hypothesis-driven feature engineering.

**RESEARCH IN CONTEXT:**

1. Systematic review: Previous attempts to predict incident dementia have relied on extensive clinical evaluations, cognitive testing, laboratory testing, neuro-imaging, genetic factors, demographics, and lifestyle variables. Applying machine learning to a large administrative claims dataset to identify individuals at increased likelihood for near-term diagnosis of dementia had not been tested.
2. Interpretation: A 50-variable model to identify those at risk for near-term diagnosis of dementia was created and validated. Based on AUC analysis, the model compared favorably with other historical attempts at modeling more traditional forms of data.
3. Future direction: Models, such as the one developed here, could be used to identify populations of higher prior probability for near-term diagnosis of dementia. These could then be subjected to more in-depth scrutiny for intervention or dementia-related research eligibility.

## 1. INTRODUCTION

As many as 35.6 million people lived with dementia worldwide in 2010 and those numbers are expected to double every 20 years to 115.4 million by 2050 (1). Within the United States, the annual number of incident cases is expected to more than double from 377,000 in 1995 to 959,000 yearly by 2050 (2) leading to a prevalence of 13.8 million (3). The total health care and long-term care costs associated with dementia are expected to reach a historic high of over quarter of a trillion dollars in the US in 2018 (4). Tools are needed to assist clinicians, public health workers, and epidemiologists in identifying individuals at risk for dementia and addressing this unfolding epidemic.

Although the upward trend in incidence of Alzheimer’s disease and the ravaging effects it has on the subject and caregivers is well documented, there has been very little published work building predictive models which help with identifying people with high risk prior to the onset of the disease. Barnes, et al. (5) developed a late-life dementia risk index, using logistic regression model developed on a cohort of 3,375 people. The model had a cStatistic of 0.81 for identifying people who were at risk of dementia in six years. The model used several important features such as demographics, cognitive scores, physical activity, results from magnetic resonance imaging (MRI) scans, social network behavior, etc. Exalto, et al. (6) developed a Cox proportional hazard model using 45 predictors with a cohort of 29,961 people with type 2 diabetes, using a combination of self-reported data, pharmacy fills for prescriptions, laboratory data, and hospitalization and outpatient records. The c-statistic for 10-year dementia risk on a validation cohort was reported as 0.74. While the results of both these studies are impressive, it is typically not possible to get such datasets including MRI scans, self-reported data, laboratory results, etc. on a large group of people. It is important to develop models using easily available data so that the model impact reaches a wider population. With rapid growth in the “baby boomer” population in the US and elsewhere requiring governments to spend billions of dollars on healthcare in an aging population, models need to be developed which can scale to large groups of people with available claims data, and can identify chronic conditions such as dementia before the onset of the disease. This early identification may assist in drug development and early treatment.

Machine learning algorithms have been used previously for developing predictive models on large administrative claims datasets using automatic feature selection. In a recent paper Rajkomar, et al. (7) developed a suite of models using machine learning algorithms on Electronic Health Records data for predicting tasks such as patient’s final discharge diagnosis, 30-day unplanned re-admission, etc. with a cStatistic of 0.9 and 0.76, respectively. McCoy, et al. (8) developed a predictive model trained on medical and pharmacy claims for 473,049 people to identify those at risk for type 2 diabetes. The model with 48 variables had a cStatistic of 0.808.

Machine learning algorithms have the potential to use large datasets to rapidly develop predictive models without specific selection of predictor variables, allowing for automated selection of high value predictors from very large numbers of potential inputs. Such models can use hundreds or even thousands of input variables related to dementia and rapidly generate useful information for clinicians, patients, pharmaceutical companies, payers, and policy-makers. Such techniques can identify connections within large datasets and algorithms that are beyond the ken of even expert clinicians. Early applications have included developing predictive models of who might develop dementia based on risk scores incorporating clinical characteristics, lab tests, neuro-imaging, and neuropsychological testing. Less investigated has been the use of machine learning to identify incident cases of Alzheimer’s disease and related dementias (ADRD) based on administrative claims data.

To that end, an effort was undertaken to develop and test a model which would predict incipient ADRD using machine learning and compare the performance of that model to previous models derived with traditional logistic regression techniques or based on individualized diagnostic testing.

## 2. METHODS

### 2.1 Data Set

This study utilizes data between 2001 and 2015 from the OptumLabs® Data Warehouse (OLDW), (9) which includes a national de-identified dataset of more than 125 million privately insured individuals that is geographically and racially diverse, including individuals of all ages (including Medicare Advantage beneficiaries ≥65 years old) and from all 50 states, with greatest representation in the Midwest and South U.S. Census Regions (10). OLDW provides full access to professional, facility, and outpatient prescription medication claims. Patient-identifying information was encrypted or removed from the study database prior to its release to the study investigators, such that it is compliant with HIPAA and exempt from Institutional Review Board review.

### 2.2 Definition of Outcome

The study outcome was a binary variable indicating a new diagnosis ADRD. The identification rules for diagnosed ADRD cases were developed based on past work (11), and extended in consultation with an Expert Advisory Panel consisting of clinicians and experts from academia. Individuals must have met at least one of the following criteria:

- a medical claim with ADRD diagnosis codes in any header position in an inpatient setting,
- a medical claim with an ADRD diagnosis code followed by another claim with an ADRD diagnosis code within 1 to 730 days; both claims can be in any setting, and the codes in any header position,
- a pharmacy claim for donepezil hydrochloride, galantamine hydrobromide, rivastigmine tartrate or tacrine hydrochloride, and
- a pharmacy claim for memantine hydrochloride along with a medical claim with an ADRD diagnosis code in any setting and any header position within 0 to 730 days. The confirmation with a diagnosis claim is required because memantine hydrochloride is also used as an augmentation therapy for anxiety disorders (OCD, ADHD, etc.) (12) as well as help slowing down the tolerance development to opioids (13).

The index date for individuals (cases), whose diagnosis was confirmed using the above criteria, was set using the earliest occurrence of a medical claim with an ADRD diagnosis code in any setting or a pharmacy claim for any of the above drugs. To ensure that the identified individuals have incident rather than prevalent ADRD, we required a 60-month period of continuous enrollment without any of the above diagnosis or pharmacy claims before the index date. Figure 1 shows examples of how the index date and confirmation date are labeled in different situations with relevant inpatient, outpatient and pharmacy claims.

**Figure 1.**
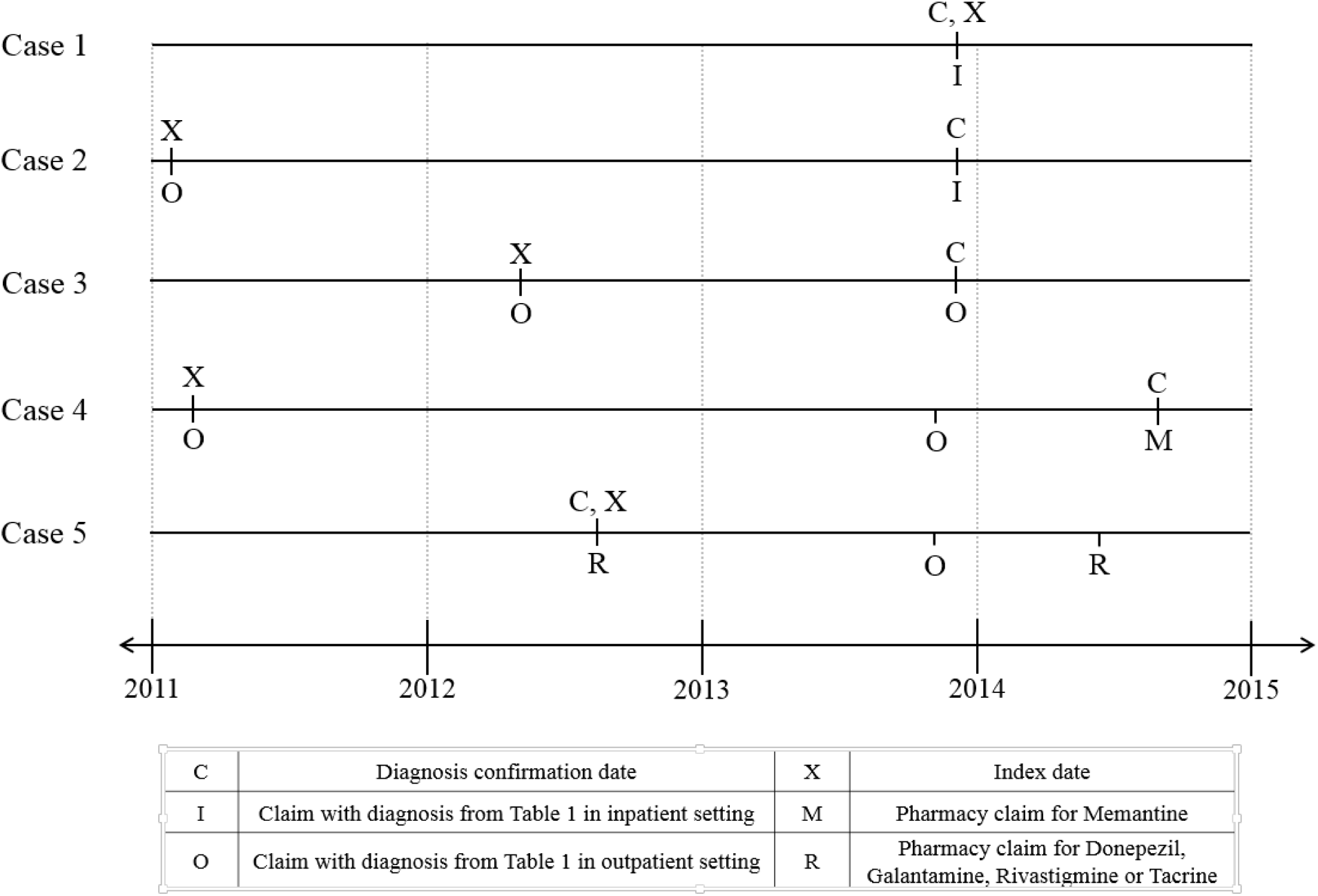
Nori *et al.*: Cases identified using different rules. Case 1 shows an individual who has an ADRD diagnosis in an inpatient setting and no previous relevant claims. So, the confirmation and index date are on the day of that claim. Case 2 has the same inpatient diagnosis claim and a previous claim in outpatient setting. So, the previous claim is used for as the index date, although it is over 730days prior to the claim in the inpatient setting. Case 3 has two claims in outpatient settings; the second claim is used as the confirmation and the first is used as the index date. Case 4 has a pharmacy claims for Memantine Hydrochloride and a diagnosis claim in an outpatient setting within 730 days. This case has a previous diagnosis claim in an outpatient setting which is used as the index date. Case 5 has multiple claims for Donepezil, Galantamin, Rivastigmin or Tacrin and the earliest of those is used for the confirmation and index dates.

### 2.3 Training and Testing

The ADRD predictive model was trained and tested using a nested case-control study design (8). Step-wise demonstration of how the study population was assembled is depicted in Figure 2. OLDW contains 59,748,354 unique individuals with any period of medical and pharmacy coverage between 1/1/2007 and 6/30/2015. The end date was chosen so that the entire study could be done using medical claims before ICD-10 diagnosis codes were required. The start date was chosen to limit the population to a size amenable for processing on the available hardware. The final cohort size (over 200,000 training observations and nearly 600,000 in test) indicated that an earlier start date was not needed to have sufficient data.

**Figure 2.**
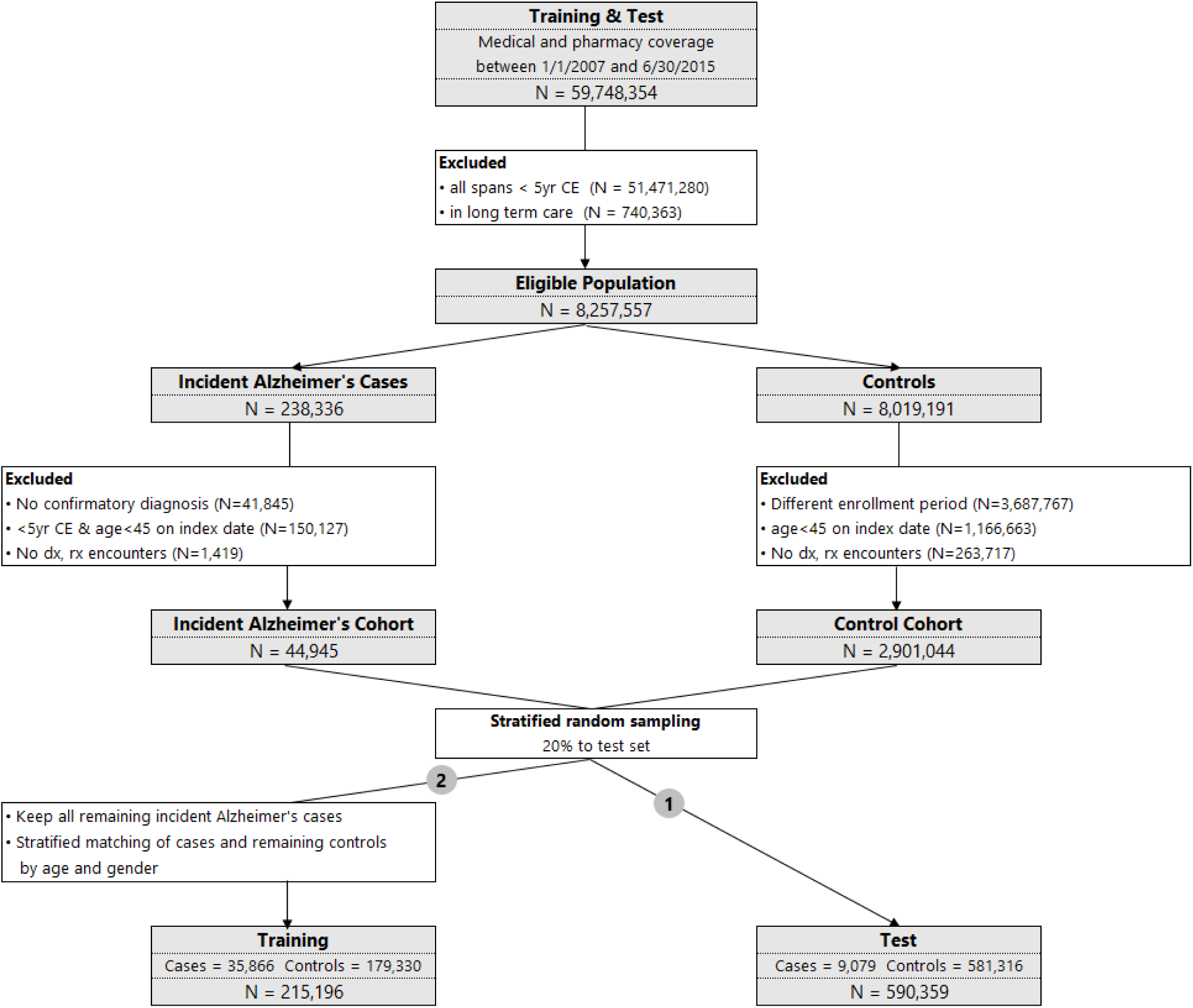
Nori *et al.*: Study design with training and validation cohorts.

We excluded 51,471,280 individuals who did not have at least one timespan with more than 5 years of continuous enrollment. Also, 740,363 individuals who are in long-term care facilities (LTC) were excluded, because LTC residence is strongly associated with pre-existing dementia (14) and also because dementia in LTC settings may be under-coded, leading the claims based identification algorithms to mark them incorrectly as cases or controls [ (15), (16)]. This resulted in 8,257,557 eligible individuals of whom 238,336 individuals had a diagnosis of ADRD based on the outcome definitions described above.

In that ADRD cohort, we excluded 41,845 individuals who did not have a confirmatory diagnosis. We also excluded 150,127 individuals who had less than 5 years of continuous enrollment immediately prior to the index date within the current enrollment span or who were less than 45 years old on the index date. To minimize bias from inclusion of individuals who had no encounters with the health care system, we excluded 1,419 people with no claims. These individuals may not be engaged in their health care management and would be poor participants in randomized controlled trials (RCT). Ultimately, there were 44,945 people in the incident ADRD cohort.

In order to minimize the risk of confounding based on duration of enrollment, the index dates for the controls cohort were selected to match the distribution of the case’s enrollment duration (Prince *et al.*, 2013). Specifically, for each person in the case cohort, we computed the time from enrollment to the index date, and divided them in percentiles 0% (1,827 days), 1% (1,832 days), …, 99% (2,719 days), 100% (2,735 days). Each person in the eligible controls population was randomly assigned one of these enrollment durations. This enrollment duration was added to their span start date to compute the index date for the controls. Out of the 8,019,191 people, these index dates for 3,687,767 individuals were outside their coverage dates and hence they were dropped from the control cohort. Another 1,166,663 people were dropped because they were less than 45 years old on the index date. Additionally, 263,717 individuals were dropped from the controls since they had no claims. Finally, there were 2,901,044 people in the control cohort.

To assemble a test population, drawing from the incident ADRD and control cohorts, we randomly selected 30% of the available cases and enrollment-duration matched controls (17); this yielded a population 9,079 cases and 581,316 controls (total 590,359). The above counts show that the original case/control cohort has about 1.5% cases. These type of two-class datasets (cases and controls) where one class constitutes a small minority of the entire data is called an imbalanced dataset (18). Machine learning algorithms, like the one used in this work, do not accurately measure model performance when faced with unbalanced datasets (19). They tend to predict the majority class data and the features in the minority class are treated as noise leading to mis-classification of the cases. The imbalanced dataset issue is handled by using a sample of the controls rather than all the controls. Such methods have been used previously in other areas as reported by Brown & Mues (20). King & Zeng (21) have presented a method for adjusting the weights in a logistic regression when one of the classes have been sampled.

The remaining 35,866 people within the ADRD incident cohort were assigned to the training population and were matched on age and gender along with enrollment duration to the 2,319,728 controls who are not in the test set so that each case could have up to 5 matched controls. The training dataset includes 215,196 individuals (35,866 cases and 179,330 controls). Thus the age/gender matching and selection of 5 matched controls implements an undersampling of the control class as the references cited above desire.

### 2.4 Independent Variables

For training and test cohorts, we collected claims for individuals for the fourth and fifth year prior to the index date. While using just the fifth year would have resulted in a model which predicts four years prior to the onset of the disease, using just one year of claims data would not capture as much data on subjects with 13 months or more between encounters. Using claims in the fourth and fifth prior years leads to a dataset with more medical data per subject. Even with two years of data per subject in the model after feature selection, 90% of individuals had 4 or fewer unique medical codes. The claims included diagnoses (ICD-9 codes), Hierarchical Ingredient Code List (HICL) drug names, procedures (CPT codes for radiology, Abraham, et al. (22)), and demographics (age and gender).

The diagnosis, drug and procedure codes were modeled using binary variables, with the value set to 1 if there was at least a single claim with a particular code. Age was modeled in ranges of 5 year increments from 40-44 to 85-89. Because of privacy concerns due to small numbers, ages greater than 89 years were mapped to 89 years. Using this process, the training model matrix had 10,363 clinical, pharmaceutical, and demographic variables (all binary) and 215,196 rows.

### 2.5 Analytic Methods

We divided the analysis into two conceptual phases: a first phase that performs feature selection and a second phase that uses the best features to create a final model. In the first phase a Lasso logistic regression algorithm was run to identify the top 50 important predictors of the dependent variable. The Lasso algorithm outperforms other machine learning algorithms such as Random Forests and Regression Trees for variable selection (23). Sensitivity tests were performed to show that using up to 500 variables did not demonstrably change the accuracy of the model.

The Lasso algorithm works by simultaneously maximizing the likelihood function (fitting the data well) and minimizing the sum of the absolute value of the coefficients (choosing a small set of features). The tradeoff between fitting the data and the number of features is managed with a regularization parameter, lambda, weighting the sum of the coefficient’s absolute values. Setting lambda very high results in no feature selection – predicting only using the prevalence of the outcome – and gradually smaller lambda values allow additional features into the model. To evaluate which value of lambda works best, 4-way cross-validation was enabled so that each fit with n variables was evaluated against 3 overlapping data sets. A full regularization path is computed starting with highest values of lambda down to lowest values of lambda on a log scale (24). The search for optimal lambda is an efficient approach to handle wide datasets with several features since it helps filter the noise and retain features with high predictive power. Since objective function used by the Lasso algorithm is a regularized version of Logistic Regression, the final list of features selected by Lasso will be a good set of predictors for the Logistic Regression algorithm. A final fit using logistic regression was used to fit all the training data (removing the cross-validation) and remove the influence of lambda on the 50 final coefficients. The Lasso algorithm was run by applying the glm function (25) in the h2o package (R version 3.5.0) which was called with parameters “nfolds=4, lambda_search=TRUE and max_active_predictors=50.” Logistic Regression was trained using the lrm function (a logistic regression algorithm) from the rms package (26).

The 50-variable ADRD model (Table 3) was used to compute scores for each individual in the training and test datasets. Scores need to be converted into a threshold above which an action will be taken with the individual. A common method to choose the threshold is to set it so the fraction of scores above the threshold is equal to the case prevalence in the population. Using this method, the amount of outreach effort that would be expended toward intervening on a potentially “at risk” group is proportional to the prevalence. Scores which were at or above the threshold were classified as at-risk for ADRD, while scores which were below the threshold were classified as not at-risk. The sensitivity, specificity and lift of the model at this threshold for training and test populations were also computed. Lift is defined here as the positive predictive value of this model divided by the case prevalence. Thus if the lift is 10, then the model reaches 10 times as many true positives as a random outreach effort.

Because the prevalence and biological causes of dementia vary based on age, the classification into predicted outcome was repeated after stratifying the individual scores using three age ranges *viz.*, 40-64, 65-74 and 75-89, and computing thresholds for each range based on the prevalence of cases in that range. The sensitivity, specificity and lift for each of the age ranges, as well as for an entire cohort using three thresholds were computed.

## 3. RESULTS

### 3.1 Cohort Characteristics

Baseline demographic and clinical characteristics of the training and test study population for cases and controls are summarized in Table 1. The age and gender-matched training population was on average 77.24 years old (SD 6.95) for cases and controls. The cases for the unmatched test set had a similar average age of 77.19 (SD 6.99), the unmatched controls were younger with an average age of 58.71 (SD 11.35). Figure 3 shows the number of all codes in the model matrix versus the cumulative percentage of observations separately for cases and controls. This figure helps to illustrate the very sparse nature of the dataset because over 55% of the cases and 80% of the controls have fewer than 3 codes (if a case or control has 3 codes, then there will be 3 ones in the row for that observation and rest zeros). This shows that three years prior to the index data, there are only a few codes in claims for majority of the members in the cohort. The data sparsity makes this a very hard binary classification problem as evidenced further in the results below.

**Table 1.** Nori *et al.*: Demographic and clinical profiles of cohorts.

|  | Training |  | Validation |  |
| --- | --- | --- | --- | --- |
|  | Cases<br>(N=35,866) | Controls<br>(N=179,330) | Cases<br>(N=9,079) | Controls<br>(N=581,316) |
| Age, years, mean (SD) | 77.24 (6.95) | 77.24 (6.95) | 77.19 (6.99) | 58.71 (11.35) |
| Gender, male, N (%) | 13,794 (38.46) | 68,970 (38.46) | 3,432 (37.80) | 267,801 (46.07) |
| Diagnosis Codes, N(%) |  |  |  |  |
| HYPERTENSION (401.9) | 21,580 (60.17) | 98,672 (55.02) | 5,419 (59.69) | 187,144 (32.19) |
| HYPERLIPIDEMIA (272.4) | 17,229 (48.04) | 82,314 (45.90) | 4,357 (47.99) | 191,833 (33.00) |
| HYPERCHOLESTEROLEMIA (272.0) | 10,768 (30.02) | 58,244 (32.48) | 2,814 (30.99) | 116,730 (20.08) |
| PAIN IN SOFT TISSUES OF LIMB (729.5) | 10,112 (28.19) | 42,702 (23.81) | 2,566 (28.26) | 91,039 (15.66) |
| OTHER MALAISE AND FATIGUE (780.79) | 9,540 (26.60) | 34,465 (19.22) | 2,403 (26.47) | 87,362 (15.03) |
| Procedure Codes N (%) |  |  |  |  |
| RADIOLOGIC EXAMINATION, CHEST (71020) | 13,915 (38.80) | 61,303 (34.18) | 3,518 (38.75) | 14,3310 (24.65) |
| DIAGNOSTIC RADIOGRAPHIC PROC (76499) | 12,759 (35.57) | 64,037 (35.71) | 3,235 (35.63) | 205,253 (35.31) |
| SCREENING MAMMOGRAPHY (77052) | 8,884 (24.77) | 51,472 (28.70) | 2,287 (25.19) | 171,086 (29.43) |
| BONE DENSITY STUDY (77080) | 6,815 (19.00) | 38,011 (21.20) | 1,746 (19.23) | 85,077 (14.64) |
| CT, HEAD OR BRAIN (70450) | 6,266 (17.47) | 18,047 (10.06) | 1,609 (17.72) | 28,423 (4.89) |
| Drug Codes N (%) |  |  |  |  |
| HYDROCODONE BIT/ACETAMINOPHEN | 4,198 (11.70) | 12,686 (7.07) | 1,046 (11.52) | 72,548 (12.48) |
| SIMVASTATIN | 3,548 (9.89) | 12,584 (7.02) | 968 (10.66) | 37,149 (6.39) |
| AZITHROMYCIN | 3,221 (8.98) | 12,561 (7.00) | 823 (9.06) | 69,553 (11.96) |
| LISINOPRIL | 3,211 (8.95) | 10,926 (6.09) | 824 (9.08) | 31,572 (5.43) |
| LEVOTHYROXINE SODIUM | 2,663 (7.42) | 9,623 (5.37) | 682 (7.51) | 29,602 (5.09) |

**Figure 3.**
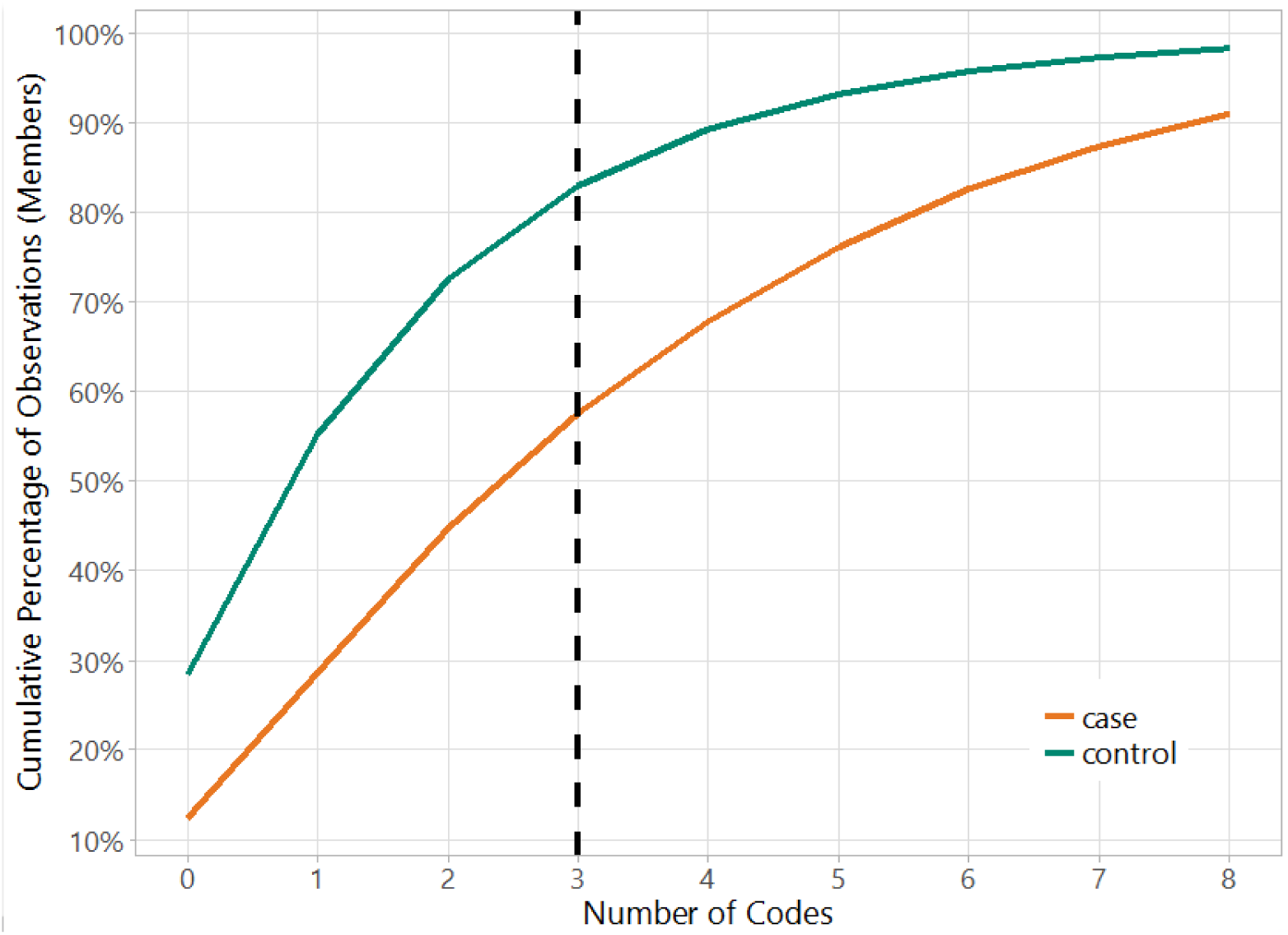
Nori *et al.*: Data Sparsity in the Cohort. Over 55% of the cases and 80% of the controls have fewer than 3 codes.

### 3.2 Model Characteristics

The 50-variable ADRD model had a mean AUC (area under the receiver operating characteristic curve) of 64.26% within [63.88%, 64.58%] for 4-fold cross-validation, and 64.25% when re-fit to the entire training population. The AUC for the test data, which was matched only on enrollment duration and not on age and gender, was 69.3%. (The improved AUC on the test data is expected because the test data set is not age matched, making the test data set easier to predict after training on age-matched data.)

Table 2 shows the sensitivity and specificity of the model for when a single threshold was used for the entire training or test cohort. The threshold for the training set is 0.20 and is computed using the 16.67% prevalence of cases due to the 1:5 matching. The sensitivity, specificity and lift are 31.9%, 86.4% and 1.9, respectively. Using the lower prevalence of 1.54% in the unmatched test cohort, a new threshold of 0.37 is computed and the same metrics are now computed as 9.9%, 98.6% and 6.4, respectively. In this cohort, the lift indicates the model reaches 6.4 times more true cases than outreach to a similarly sized random group.

**Table 2.** Nori *et al.*: Model sensitivity and specificity computed using a single threshold for the entire cohort based on the prevalence of the cases and for each age range based on the prevalence of the cases in those age ranges. Age and gender matching in training yields different prevalence and measures.

| **Cohort** | **Age** | **Prevalence** | **Threshold** | **Sensitivity** | **Specificity** | **Lift** |
| --- | --- | --- | --- | --- | --- | --- |
| Training | all | 16.67% | 0.20 | 31.9% | 86.4% | 1.9 |
|  | 15-64 | 16.67% | 0.18 | 41.8% | 88.4% | 2.5 |
|  | 65-74 | 16.67% | 0.19 | 39.1% | 87.8% | 2.3 |
|  | 75-99 | 16.67% | 0.21 | 29.6% | 85.9% | 1.8 |
|  | computed | 16.67% |  | 32.1% | 86.4% | 1.9 |
| Test | all | 1.54% | 0.37 | 9.9% | 98.6% | 6.4 |
|  | 15-64 | 0.14% | 0.61 | 3.3% | 99.9% | 23.3 |
|  | 65-74 | 1.77% | 0.40 | 9.8% | 98.4% | 5.6 |
|  | 75-99 | 8.46% | 0.26 | 19.2% | 92.5% | 2.3 |
|  | computed | 1.54% |  | 16.4% | 98.7% | 10.7 |

To enhance the model predictions, we chose thresholds that varied based on age-based prevalence. The performance can be computed as shown in Table 2. The table shows that the sensitivity increases with age for the test cohort. The sensitivity (specificity) for the test cohort increases from 9.9% (98.6%) to 16.4% (98.7%) by using the age-based thresholds. Thus the age specific thresholds increase the sensitivity 64% without a demonstrable change in the specificity.

Figure 4 shows a density plot of the distribution of scores for the cases and controls in the training and test cohorts, for the different groups of ages and for all ages. The dashed line indicates the threshold; all scores greater than or equal (lesser than) that line would be classified as cases (controls). These plots depict separation of the cases and controls at higher score range, with a large overlap in scores at the lower range.

**Figure 4.**
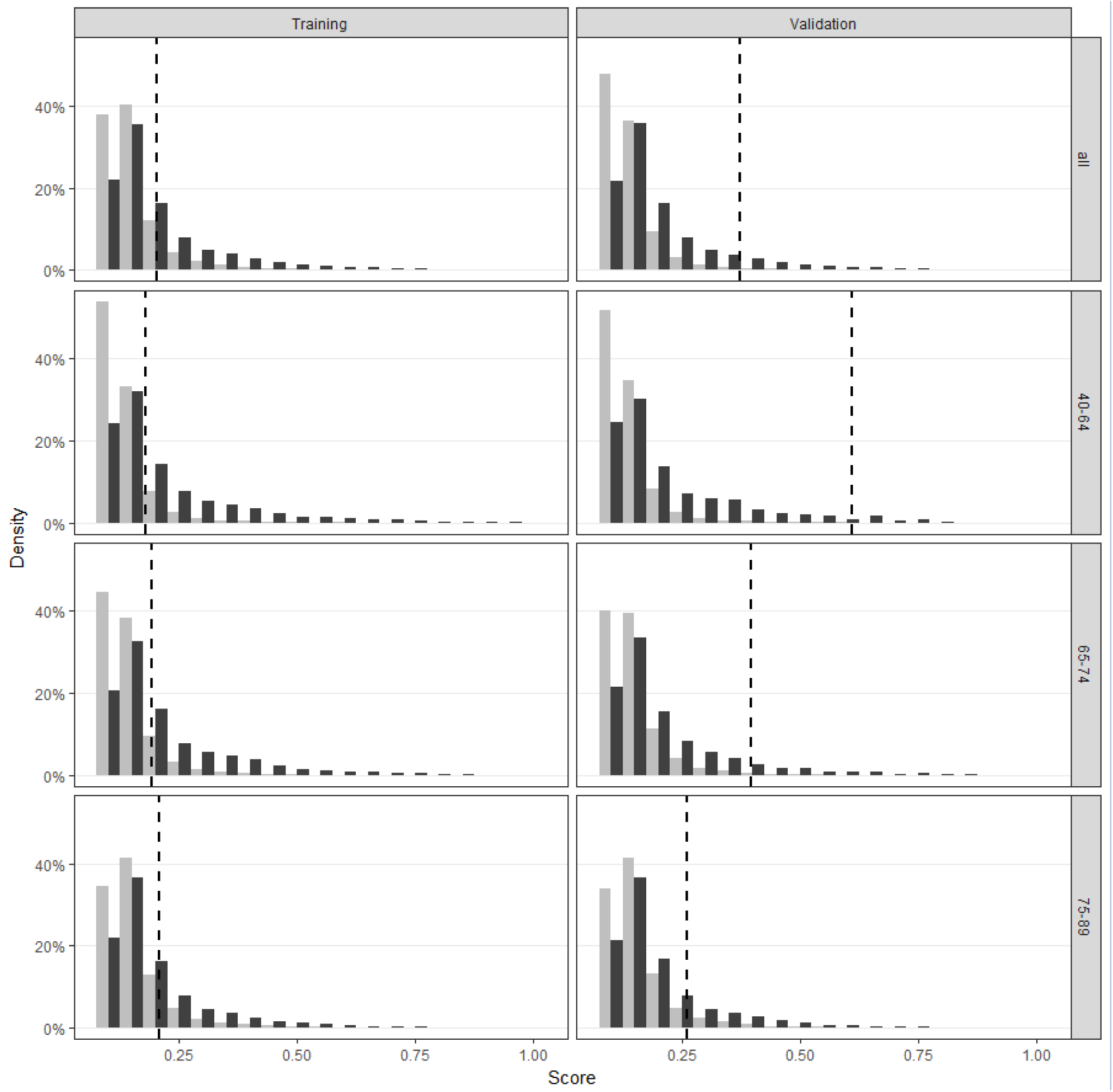
Nori *et al.*: Distribution of scores for cases and controls for different age ranges, demonstrating substantial overlap in scoring between cases and controls. The vertical lines show a proposed cut point for classification.

Table 3 shows the coefficients, significance and variance inflation factor (VIF) for each variable in the model. The intercept term is the model constant representing the default risk for a male patient aged 65-69 with no other features. The maximum VIF for any variable is 1.46 indicating that the model has minimal collinearity. The top four diagnosis codes with a coefficient greater than 1 are Memory loss (780.93), Parkinson ‘s disease (332.0), Mild Cognitive Impairment (331.83) and Bipolar Disorder (296.80). The top 5 pharmacy codes are psychoactive drugs.

**Table 3.** Nori *et al.*: Clinical diagnosis, procedure and pharmacy variables included in the model, coefficients and variance inflation factors (VIF). The intercept has no VIF because it is a constant and does not vary across the observations due to age/gender matching. The specific ICD-9 codes for diagnosis and CPT-4 codes for procedures used to identify these variables are shown in parenthesis.

| Type | Variable Description | Coefficient | Pr(>\|Z\|) | VIF |
| --- | --- | --- | --- | --- |
|  | INTERCEPT (MALE, 65-69) | -1.96 | <0.0001 |  |
| ICD-9-CM | MEMORY LOSS (780.93) | 1.33 | <0.0001 | 1.02 |
|  | PARALYSIS AGITANS (332.0) | 1.17 | <0.0001 | 1.01 |
|  | MILD COGNITIVE IMPAIRMENT, SO STATED (331.83) | 1.14 | <0.0001 | 1.01 |
|  | BIPOLAR DISORDER, UNSPECIFIED (296.80) | 1.00 | <0.0001 | 1.01 |
|  | UNSPECIFIED PSYCHOSIS (298.9) | 0.44 | <0.0001 | 1.06 |
|  | LOSS OF WEIGHT (783.21) | 0.42 | <0.0001 | 1.02 |
|  | DEPRESSIVE DISORDER, NOT ELSEWHERE CLASSIFIED (311) | 0.37 | <0.0001 | 1.10 |
|  | ALTERED MENTAL STATUS (780.97) | 0.33 | <0.0001 | 1.16 |
|  | PERSONAL HISTORY OF FALL (V15.88) | 0.32 | <0.0001 | 1.05 |
|  | OTHER CONVULSIONS (780.39) | 0.32 | <0.0001 | 1.04 |
|  | UNSPECIFIED FALL (E888.9) | 0.27 | <0.0001 | 1.06 |
|  | OTHER CHRONIC PAIN (338.29) | 0.28 | <0.0001 | 1.03 |
|  | ACUTE, BUT ILL-DEFINED, CEREBROVASCULAR DISEASE (436) | 0.24 | <0.0001 | 1.33 |
|  | URGE INCONTINENCE (788.31) | 0.24 | <0.0001 | 1.07 |
|  | OTHER ALTERATION OF CONSCIOUSNESS (780.09) | 0.23 | <0.0001 | 1.09 |
|  | UNSPECIFIED CONSTIPATION (564.00) | 0.21 | <0.0001 | 1.05 |
|  | UNSPECIFIED URINARY INCONTINENCE (788.30) | 0.21 | <0.0001 | 1.08 |
|  | ENCOUNTER FOR LONG-TERM (CURRENT) USE OF OTHER MEDICATIONS (V58.69) | 0.17 | <0.0001 | 1.03 |
|  | LACK OF COORDINATION (781.3) | 0.15 | <0.0001 | 1.09 |
|  | OTHER MALAISE AND FATIGUE (780.79) | 0.11 | <0.0001 | 1.15 |
|  | DIABETES MELLITUS (250.02) | 0.12 | <0.0001 | 1.30 |
|  | ABNORMALITY OF GAIT (781.2) | 0.10 | <0.0001 | 1.20 |
|  | DIZZINESS AND GIDDINESS (780.4) | 0.11 | <0.0001 | 1.13 |
|  | UNSPECIFIED CEREBRAL ARTERY OCCLUSION WITH CEREBRAL INFARCTION (434.91) | 0.11 | 0.0083 | 1.35 |
|  | DIABETES MELLITUS (250.00) | 0.07 | <0.0001 | 1.42 |
|  | EDEMA (782.3) | 0.07 | <0.0001 | 1.10 |
|  | MUSCLE WEAKNESS (GENERALIZED) (728.87) | 0.06 | 0.0277 | 1.14 |
|  | URINARY TRACT INFECTION, SITE NOT SPECIFIED (599.0) | 0.03 | 0.0526 | 1.12 |
| CPT | SCREENING MAMMOGRAPHY; COMPUTER-AIDED DETECTION (77052) | -0.25 | <0.0001 | 1.03 |
|  | COMPUTED TOMOGRAPHY, HEAD OR BRAIN (70450) | 0.16 | <0.0001 | 1.42 |
|  | RADIOLOGIC EXAMINATION, CHEST; SINGLE VIEW, FRONTAL (71010) | 0.05 | 0.0011 | 1.28 |
| Medication (HICL Descripti) | VENLAFAXINE HCL | 0.52 | <0.0001 | 1.02 |
|  | DULOXETINE HCL | 0.45 | <0.0001 | 1.05 |
|  | TOLTERODINE TARTRATE | 0.32 | <0.0001 | 1.07 |
|  | SERTRALINE HCL | 0.29 | <0.0001 | 1.05 |
|  | CITALOPRAM HYDROBROMIDE | 0.25 | <0.0001 | 1.06 |
|  | POTASSIUM CHLORIDE | 0.24 | <0.0001 | 1.35 |
|  | OXYBUTYNIN CHLORIDE | 0.22 | 0.0001 | 1.08 |
|  | HYDROCODONE BIT/ACETAMINOPHEN | 0.19 | <0.0001 | 1.27 |
|  | PROPOXYPHENE/ACETAMINOPHEN | 0.15 | <0.0001 | 1.09 |
|  | SULFAMETHOXAZOLE/TRIMETHOPRIM | 0.15 | <0.0001 | 1.12 |
|  | METFORMIN HCL | 0.13 | 0.0001 | 1.40 |
|  | BLOOD SUGAR DIAGNOSTIC | 0.12 | 0.0011 | 1.38 |
|  | LISINOPRIL | 0.11 | <0.0001 | 1.17 |
|  | CEPHALEXIN MONOHYDRATE | 0.10 | 0.0003 | 1.12 |
|  | SIMVASTATIN | 0.10 | <0.0001 | 1.17 |
|  | CLOPIDOGREL BISULFATE | 0.09 | 0.0054 | 1.11 |
|  | TRAMADOL HCL | 0.09 | 0.0097 | 1.13 |
|  | GABAPENTIN | 0.07 | 0.0914 | 1.12 |
|  | FUROSEMIDE | 0.00 | 0.9396 | 1.46 |

## 4. DISCUSSION

A recent systematic review of dementia risk prediction models (27) found models that could be grouped into five categories: (1) demographic factors only; (2) cognitive based (cognitive test scores); (3) health variables and risk factors; (4) genetic risk scores; and (5) multi-variable models which combined demographic with health and lifestyle factors. Previously, machine learning techniques have been primarily being applied to clinical data to successfully identify early cases of ADRD (28), to cluster patients into fast versus slow progression sub-types (29), to distinguish mild cognitive impairment or normal aging from early dementia [ (28), (30)], and to assist in the interpretation and clinical significance of findings from neuro-imaging studies [ (31), (32), (33), (34), (35) (36)]. Performing such work using administrative claims data may offer a larger pool for analyses and identification, since such claims are more widely available for large populations.

As noted earlier, a national de-identified dataset was used to train and test the models. While the census region-level information for members in the cohort is available, it was not used to control for potential regional differences in the model for the following reasons. First is that while it may help with adjusting for access to care, it will not help with addressing differences in coding behavior between providers, difference in insurance plan types, etc. Second, we wanted the model to be usable in a variety of production settings with claims coming in from different sources without the need for that additional information. This dataset has a combination of so many benefit plans, regions, etc. potential bias in any one of them would not effect the overall model.

The performance of this claims-based model with an AUC of 64% compares favorably with other models based more directly on clinical data collected about risk factors and health variables. The inclusion of a validation component in this study represents a refinement not often found in previous studies. The aforementioned meta-analysis of 21 papers on dementia risk prediction found only 4 with validation components (27). AUC values in these papers ranged from 49% to 78%. These models used various non-claims based variables including genotype. These extra variables come at high cost and based on those published AUC values, do not always lead to better models.

A review of the medications and diagnosis codes that remained in the final model (Table 3) suggests that helpful inferences can be drawn from machine-learning derived predictors. For example, six variables (12% of the model) related to falls, dizziness, gait disorders, or weakness, suggesting a subacute phase of progression. Additionally, diagnoses and medications related to vascular disease and to diabetes mellitus were represented, which supports clinical data suggesting overlap of risk factors for cardiovascular disease with vascular dementia [ (37), (38)] and an association of dementia risk with diabetes mellitus [ (39), (40), (41)]. Psychiatric symptoms and psychoactive medications were also unsurprisingly present. Screening mammography proved to be the only code with a negative coefficient indicative of a protective effect. Because the application of screening procedures can be tempered by clinical judgment which incorporates such factors as life expectancy and quality of life, the presence of this screening test might be a marker for healthier and less impaired individuals who would therefore be at lower risk for developing dementia.

This model becomes increasingly useful as potential disease-modifying treatments for dementia are developed to a stage for clinical testing. Thus, the ability to achieve a lift of 6.4 means that a patient identified by the model will be 6.4 times more likely to be diagnosed in the near-term with dementia. An identified cohort with such enhanced prior probability could be much more cost-effectively screened for clinical research than an unselected population.

A limitation of our study is that dementia may be undercoded, presenting a challenge for training models; in one study, Alzheimer’s disease and related dementias was recorded as a diagnosis for less than 25% of patients with moderate to severe cognitive impairment (42); and in another, physicians were unaware of cognitive impairment in more than 40% of their cognitively impaired patients (43). Among participants in a Medicare Alzheimer’s Disease Demonstration, less than 20% of participants were classified with dementia of the Alzheimer type based on a year’s worth of claims data, although 68% carried that diagnosis upon referral (44). A review of seven studies examining the extent to which dementia is omitted as a cause of death, found that the reporting on death certificates ranged from a 7.2% to 41.8% (45).

